# Desmosomal coupling in apoptotic cell extrusion

**DOI:** 10.1101/816280

**Authors:** Minnah Thomas, Benoit Ladoux, Yusuke Toyama

## Abstract

The mechanical coupling of epithelia enables coordination of tissue functions and collective tissue movements during different developmental and physiological processes. This coupling is ensured by cell-cell junctions, including adherens junctions (AJs) and desmosomal junctions (DJs) [1, 2]. During apoptosis, or programmed cell death, a dead cell is expelled from the tissue by coordinated processes between the dying cell and its neighbors. Apoptotic cell extrusion is driven by actomyosin cable formation and its contraction, and lamellipodial crawling of the neighboring cells (Fig. S1A-A’’, Movie S1) [3–6]. Throughout cell extrusion, the mechanical coupling of epithelia needs to be maintained in order to preserve tissue homeostasis [3]. Although much is known about the regulation of AJs in apoptotic cell extrusion [6–9], the role and dynamics of DJs during this process remains poorly understood. Here, we show that DJs stay intact throughout and are crucial for apoptotic cell extrusion. Pre-existing DJs between the apoptotic cell and neighboring non-dying cells remain intact even during the formation of *de novo* DJs between non-dying cells, suggesting that the neighboring cells possess two DJs in the middle of apoptotic cell extrusion. We further found that an actomyosin cable formed in the vicinity of DJs upon apoptosis, and subsequently deviated from DJs during its constriction. Interestingly, the departure of the actomyosin cable from DJs coincided with the timing when DJs lost their straightness, suggesting a release of junctional tension at DJs, and a mechanical coupling between DJs and actomyosin contractility. The depletion of desmoplakin, which links desmosomes and intermediate filaments, resulted in defective apical contraction and an inability to form *de novo* DJs, leading to a failure of apoptotic cell extrusion. Our study provides a framework to explain how desmosomes play pivotal roles in maintaining epithelial sheet integrity during apoptotic cell extrusion.

## RESULTS AND DISCUSSION

During apoptotic cell extrusion, adherens junction components including E-cadherin and α- and β-catenins display a reduction in their levels at the interface between the apoptotic cell and neighboring cells. This reduction coincides with membrane disengagement, actomyosin cable formation, and relaxation of tissue tension [6]. Here, we investigate the dynamics of desmosomal junctions (DJs), which are a part of the tripartite epithelial junctions known to influence dynamic processes such as collective cell migration [10] and skin differentiation [11, 12], during cell extrusion.

### Desmosomal junctions stay intact throughout apoptotic cell extrusion

To understand how DJs were remodeled during the course of apoptotic cell extrusion, we used a UV laser to induce DNA damage and subsequent apoptotic cell extrusion (Supplemental experimental procedures, [5]), and followed the progression of extrusion using confocal microscopy (Fig. 1A). We first examined the distribution of desmoglein, one of the two cadherin types found in DJs, by expressing EGFP-tagged desmoglein 2 in wild-type (WT) Madin-Darby canine kidney (MDCK) cells (Fig. 1B, S1B-B’). We took advantage of the non-homogeneous expression of desmoglein 2-FLAG-EGFP within a tissue, and induced apoptosis in a non-EGFP expressing cell that was adjacent to EGFP positive cells in order to follow the changes in desmoglein in the neighboring non-dying cells. Stills from time-lapse movies showed that in contrast to AJs, (Fig. S1C-C’ and [6]) DJs between apoptotic and neighboring non-dying cells did not show a reduction in desmoglein 2 levels throughout extrusion (red arrows in Fig. 1B, Movie S2), but rather exhibited up to a 2.2±0.5 fold increase in intensity (Supplemental experimental procedures, Fig. 1B’). Furthermore, neighboring non-apoptotic cells formed punctate *de novo* DJs at the basal section of the cell, once lamellipodia formed in neighboring cells apposed to each other (blue arrows in Fig. 1B, S1D). This was seen more clearly in the transverse view (Fig 1C), which shows a pre-existing DJ located at the lateral section of the cells (red arrows), and a nascent DJ with the new neighboring cell, formed at the tip of lamellipodia (blue arrows). Later, these *de novo* DJs became matured junctional plaques that are positioned more apically on the lateral membrane, suggesting the neighboring non-dying cells form mature DJs underneath the apically extruding apoptotic cell. We further noticed that DJs lost their straightness at later stages of cell extrusion (red arrows in Fig. 1B, see also later section). A similar localization was observed in the endogenous desmosome during extrusion (Fig. S1E) and during reactive oxygen species (ROS)-induced apoptosis. (Fig. S1F-F”).

**Fig 1:**
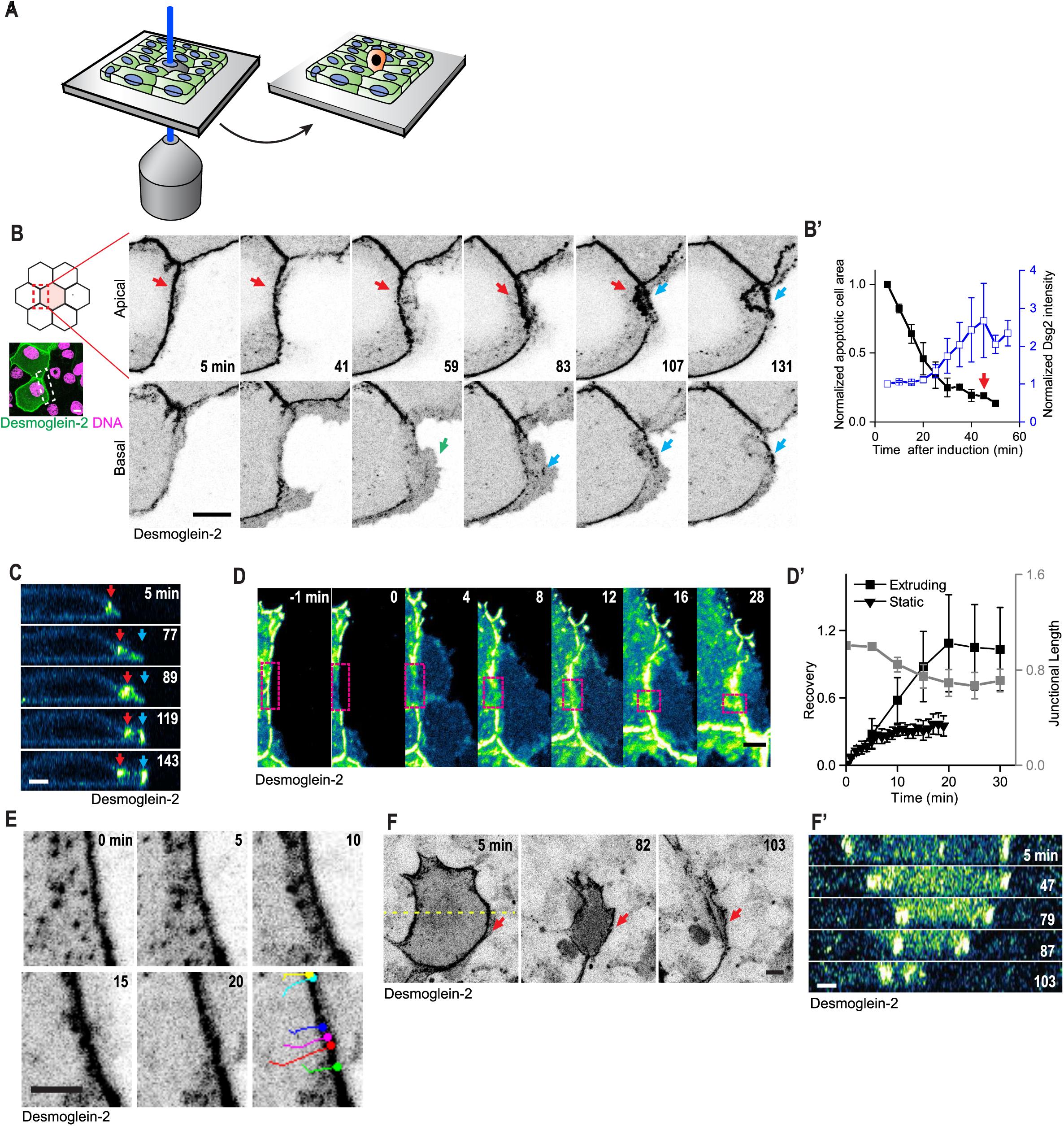
Desmosome dynamics during apoptotic cell extrusion. A) Diagram illustrating the experimental setup for targeted apoptosis induction in MDCK monolayers. B) Confocal images showing DJ dynamics at apical (top panel) and basal (bottom panel) sections of the cell. Red, blue, and green arrows indicate the pre-existing DJ, *de novo* DJ, and lamellipodia, respectively. The survey view (left) of the extrusion site containing the ROI (White dotted box) used in B. B’) Graphs plotting normalized apoptotic cell area (black) and DJ intensity (blue) during extrusion (n=6 independent experiments). Red arrow indicates average extrusion time. C) Transverse view of a neighboring cell showing DJ dynamics during extrusion. Red and blue arrows indicate pre-existing and *de novo* DJs, respectively. D) Time lapse images showing photobleaching and recovery (ROI: Magenta box) of Desmoglein 2 during extrusion. D’) Graphs plotting normalized recovery curves of Desmoglein 2 (black square) and corresponding junctional length (gray) during early apoptotic cell extrusion. Normalized recovery curve of Desmoglein 2 from non-apoptotic junction (static junction) is also shown for comparison (n=5 independent experiments each). E) Confocal images showing the movement of Desmoglein 2 puncta toward DJs during early extrusion. The bottom panel shows the tracks of Desmoglein 2 puncta over time (n=5 independent experiments). F) Confocal images showing the temporal localization of Desmoglein 2-FLAG-EGFP in the dying cell during extrusion. Red arrows indicate a junctional interface (n=5 independent experiments). F’) Transverse view of apoptotic cell extrusion generated along the yellow dotted line highlighted in F. The data represented in B’ and D’ represent mean ± SEM. Scale bars, 5 μm except F (10 μm).

To understand whether pre-existing DJs are dynamic during extrusion, we performed fluorescence recovery after photobleaching (FRAP) of EGFP-tagged desmoglein 2 expressed in non-apoptotic cells at different stages of extrusion, classed as before, early, and late (i.e., after DJs lost their straightness) stage of cell extrusion (Fig. 1D, S2A-A”). Although it is challenging to quantitatively interpret the recovery rate because of the shortening of DJs during constriction, there was turnover of desmoglein 2 at DJs (Fig. 1D’, S2A’), suggesting that pre-existing DJs are dynamic throughout apoptotic cell extrusion. This recovery and the subsequent increase in desmoglein 2 intensity during extrusion originated, in part, from the incorporation of desmoglein 2 puncta from the cytoplasm into the DJs (Fig. 1E, Movie S3, Fig. S2B-B’), which is analogous to a previous observation made during DJ formation [13]. In order to uncover whether the dynamic nature of DJs is crucial for cell extrusion, we prevented the turnover of DJs by pre-treating the tissue with GÖ6976, which is a PKC inhibitor that prevents the transformation of stable desmosomes into a ‘remodelable’ state [14] .Under such conditions, extrusion was defective, i.e., more cells failed to successfully extrude within two hours after UV irradiation (Fig. S2C-C’), indicating that desmosome remodeling is required for extrusion. Next, we examined DJs in the apoptotic cell by tracking desmoglein 2-FLAG-EGFP in a dying cell, surrounded by non-labeled neighbors. Although biochemical analysis of desmosomal cadherins has reported that they undergo both caspase- and matrix metalloprotease (MMP)-mediated cleavage [15–20], we did not observe a strong reduction in desmoglein 2 levels in the apoptotic cell throughout extrusion (Fig. 1F-F’). Together, our data showed that pre-existing DJs between apoptotic and neighboring cells remain intact even during the formation of *de novo* DJs between non-dying cells, suggesting that the neighboring cells possess two DJs in the middle of apoptotic cell extrusion. This further raises the possibility of intercellular force transduction across the interface during extrusion.

### Cytokeratin-18 in the neighboring cell aligns and accumulates during cell extrusion

Junctional remodeling is often accompanied by reorganization of the associated cytoskeleton. Therefore, we examined the dynamics of the DJ-associated intermediate filament protein cytokeratin-18 (CK18) tagged with mEmerald (Fig. 2A), during apoptotic cell extrusion. Similar to our desmoglein 2 experiments, we induced apoptosis in a non-fluorescent cell next to mEmerald-positive cells to follow the changes in cytokeratin within the neighboring non-dying cells. Stills from time-lapse movies showed that cytokeratin filaments that were linked to pre-existing DJs realigned towards the apoptotic cell during extrusion (Fig. 2A top, Movie S4-S5, Fig. S2D). We then quantified the orientation of cytokeratin filaments by measuring the structure tensor of the image [21] (Supplemental experimental procedures), which were represented as color-coded orientation plots (Fig. 2B). The orientation of the filaments in the vicinity of DJs at the interface between dying and non-dying cells was random (i.e. mixed colors) at the early stage of extrusion and became more polarized (i.e. cold colors) towards the apoptotic cell at late stage (Fig. 2B-B’). These observations were further supported by plotting the distribution of orientation angles of filaments (Fig. 2B”). Another pool of dynamic cytokeratin-18 was found within the lamellipodia formed in the neighboring cells (Fig. 2A, white arrow). Quantitative measurement of the fluorescence intensity of cytokeratin18 at the junctional interface within neighboring cells showed that the intensity increased as apoptotic cell extrusion progressed (Fig. 2A’). Similar accumulation of cytokeratin-18 was observed during extrusion in the endogenous protein (Fig. 2C), and accumulation of cytokeratin-5 and 8 have been reported in oncogenic cell extrusion [22]. To understand the kinetics of keratin reorganization in the course of extrusion, we took advantage of the photo-convertible protein, Dendra2-tagged cytokeratin-18, and performed photoconversion experiments in the neighboring cells. We found that the intensity of the red signal, which represents the photoconverted pre-existing keratin, did not change over time (Fig. S2E-E”). By contrast, the intensity of the green signal decreased upon photoconversion and increased as extrusion progressed, suggesting that new cytokeratin-18 was recruited in the vicinity of the interface between dying and non-dying cells (Fig. S2E-E”). Together, the alignment and accumulation of cytokeratin during apoptotic cell extrusion prompted us to speculate that the viscoelastic gel-like cytoplasm in neighboring cells undergoes stretching and thus remodels towards the extrusion site. To test this hypothesis, we performed laser ablation to assess the tension in keratin filaments during extrusion. The reoriented keratin filaments at the late phase of extrusion were associated with significantly higher tension in comparison to the random keratin network at the early phase of constriction as shown by the recoil velocity (0.17±0.06 µm/s vs 0.07±0.04 µm/s) (Fig. 2D-D’, Movie S6). This indicates that desmosome-coupled keratin network is stretched and DJs bear mechanical force during extrusion [23].

**Fig 2:**
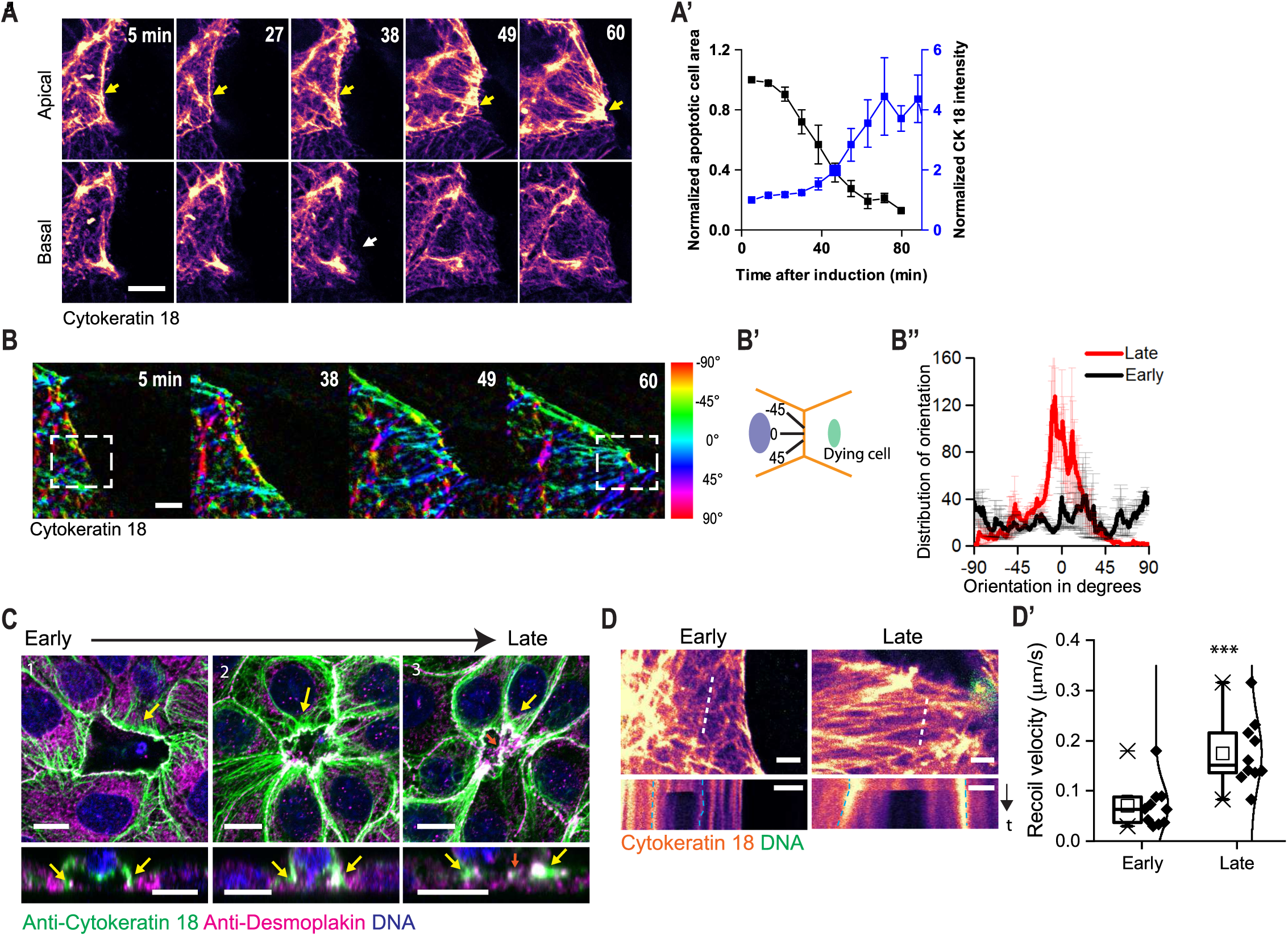
Cytokeratin-18 rearranges during apoptotic cell extrusion. A) Confocal images showing the dynamics of mEmerald-Cytokeratin 18 within the neighbouring cell at apical (top panel) and basal (bottom panel) sections of the cell during extrusion. Yellow arrows denote the interface between the apoptotic cell and the neighbouring cell. White arrow highlights Cytokeratin-18 in the lamellipodia. A’) Evolution of normalized apoptotic cell area (black) and Cytokeratin-18 intensity (blue) (n=4 independent experiments). B) Orientation map of Cytokeratin-18 network at different stages of extrusion. White dotted box denotes the region used to measure the orientation. B’) Illustration describing the filament position and orientation angle. B’’) Graph plotting the distribution of Cytokeratin-18 filament orientation at early (black) and late (red) phases of constriction (n=4 independent experiments). C) Confocal images showing localization of desmosomes (green) and Cytokeratin-18 (magenta) during different stages of extrusion. Yellow arrows highlight Cytokeratin-18 at the interface. Orange arrow shows the *de novo* junction formation (n=80 extrusion sites from 4 independent staining experiments). D) Confocal images showing keratin network at early and late stages of constriction. White dotted line shows the site of laser ablation. The kymograph shows the recoil of the filaments. Blue dotted line highlights the edge of the keratin filament tracked. D’) Recoil velocity of keratin filaments at early and late stages of constriction (n=10 independent experiments for each condition). The data represented in A’, B’’ and D’ represent mean ± SEM. Scale bars, 5 μm except C (10 μm). Unpaired two tailed t-test was performed for statistical analysis of D. ***: p<0.001.

### An actomyosin cable forms in close proximity to the DJs at the early stage, and decouples from DJs at the late stage of cell extrusion

We next assessed the relationship between DJs and apoptosis-induced actomyosin cable formation by imaging LifeAct-Ruby (marker for F-actin), and GFP-tagged desmoglein 2 (Fig. 3A-A’). Time-lapse confocal images showed that an apoptosis-associated actomyosin cable formed in close proximity to the pre-existing DJs (t=30, Fig. 3A-A’). This is consistent with a recent study that shows the recruitment of actin to desmoglein 1 at DJs, through cortactin- and Arp2/3-mediated actin polymerization during keratinocyte delamination [12]. Actin and DJs moved together during the early stage of cell extrusion, which is driven by contraction of the actomyosin cable (t=30, Fig. 3A-A’, S3A-A’)[24]. Later, the actin cable spatially deviated from the DJs and shifted towards the basal side of the apoptotic cell (Fig. 3A-A’, Movie S7, Fig. S3B). Transverse views and quantification of the z-positions of the DJs and the actomyosin cable at different stages of cell extrusion further supported our observations (Fig. 3A’-A”, S3C). Consistent with the analyses of DJs and actin, cytokeratin-18 linked to the DJs also segregated from the actin cable (Fig. S3D-D’, Movie S8). Intriguingly, the departure of the actomyosin cable from the DJs coincided with the timing when DJs lost their straightness (Fig. 3A, 1B). The quantification of DJ straightness clearly showed that they became wavier upon decoupling of the actin cable (Fig. 3B-B’), suggesting a release of junctional tension [25] at DJs. Our observations prompted us to speculate that DJs and actomyosin contractility are somehow coupled during apoptotic cell extrusion and in native cell-cell junctions outside of extrusion.

**Fig 3:**
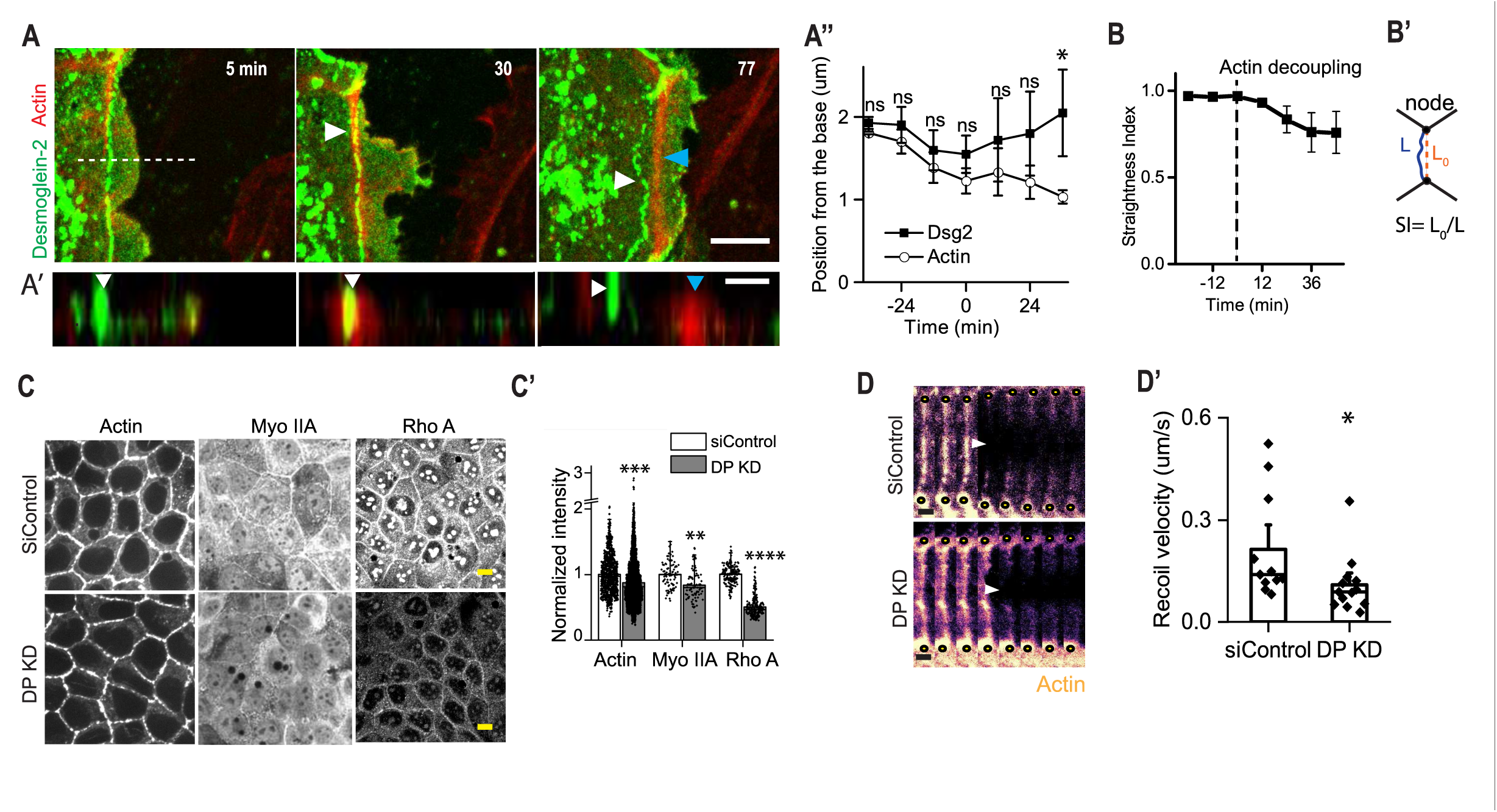
Interplay of desmosomes and actomyosin in apoptotic and steady state junctions. A) Confocal images showing the localization of actin (blue arrowhead) and desmoglein 2 (white arrowhead) at the apical section of the cell. White dotted line shows neighbour cell contours. A’) Transverse view of a neighboring non-dying cell along the white dotted line highlighted in A. Blue and white arrowheads point to actin and desmoglein 2 respectively (n=7 independent experiments). A”) Quantification of the relative positions of actin and desmoglein 2 at different phases of extrusion (n=5 independent experiments). B) The straightness index (SI= Junctional node displacement (L0)/ path length, (L)) of the junction during extrusion. The dotted line denotes the time when actin decoupled from DJs (n=5 independent experiments). B’) Illustration showing the computation of straightness index at the interface. C) Confocal images showing junctional contractile machinery including actin, myosin IIA, and RhoA in siControl and DP-depleted MDCK monolayers. C’) Graph showing the relative junctional intensities of actin, myosin IIA and RhoA in scrambled control and DP-depleted junctions. The junctional intensity of DP-depleted junctions was normalized to the average of the corresponding siControl junctions (Actin: n=861 junctions (siControl) and 1205 junctions (DP KD); MyoII: n= 141 (SiControl) and 219 (DP-KD) junctions; RhoA: n=175 (SiControl) and 211 (DP KD) junctions from 3 independent staining experiments respectively). D) Time lapse images showing actin (Lifeact-Ruby) in both siControl control and DP-depleted junctions, before and after junctional ablation respectively. The yellow dot indicates the position of the junctional node. The white arrowhead shows the site of laser ablation. D’) Recoil velocity after laser ablation at intercellular junctions in siControl control and DP-depleted monolayers (n=12 (siControl) and 16 (DP KD) from 3 independent experiments respectively). The data from A’’, B, C’ and D’ represent mean ± SEM. Paired t-test was used for statistical analysis of A’’. Unpaired two tailed t-tests were performed for statistical analysis of C’ and D’ respectively. *: p<0.05, **:p<0.01, ****: p<0.0001. Scale bars, 10 μm except A’ and D, (2 μm).

### Desmosome-depleted junctions have aberrant junctional actomyosin networks and weakened junctional tension

To test this hypothesis, we altered desmoplakin (DP) levels by using short interfering RNA (siRNA), and assessed the factors related to junctional tension in a monolayer that was not undergoing apoptosis (Fig. S3E-F). First, we found that the levels of actin, non-muscle myosin IIA, and RhoA were all lower in siRNA-mediated DP knockdown (KD) monolayers than in scrambled control (Fig. 3C-C’), suggesting that DP-depletion was associated with aberrant junctional actomyosin networks and contractility. This was further supported by measurements of the junction-to-cytoplasm ratios of myosin IIA and RhoA in control and knockdown monolayers (Fig. S3G). Our results are in agreement with the defective junctional actomyosin networks observed during the depletion of desmoglein 1 from tissue monolayers [12], indicating that depletion of desmosome components leads to an inability to retain the contractile machinery. This may arise from defects in the shuttling of myosin II between its junctional and cytoplasm pools, or in myosin II activation [26, 27]. We then performed laser ablation to characterize changes in junctional tension within the monolayer [25] (Fig. 3D, Movie S9). The initial recoil velocity after laser ablation (Supplemental experimental procedures) is a good approximation of the junctional tension before ablation [28]. DP-depleted monolayers showed a significantly lower junctional tension as compared to control monolayers (Fig. 3D-D’), which is in agreement with a recent report that demonstrated lower junctional force upon desmosome depletion [12, 27]. It is important to note that tension levels in the non-apoptotic monolayer (Fig. 3D’) are lower than the tension at the junctions between apoptotic and neighboring cells (Fig. S3H-H”, Movie S10). Together, our results show that DJs directly or indirectly influence actomyosin contractility and this in turn, affects junctional tension within a tissue. This coupling could also affect dynamic cellular processes, including apoptotic cell extrusion.

### Desmosomes are crucial for successful apoptotic cell extrusion

To further investigate the functional significance of DJs, we studied apoptotic cell extrusion in desmoplakin-depleted monolayers. LifeAct-Ruby was expressed to visualize actin distribution and cell shape (Fig. 4A). A few cases of extrusion event (5%, n=70) were associated with a tissue tearing phenotype at the interface between the apoptotic and neighboring non-dying cells (Fig. 4B). In the non-tearing event, more than half of the apoptotic cells (51%, n=70) showed defective extrusion in DP-depleted monolayers, whereas in the control tissue all cells (n=70) were successfully extruded within two hours (Fig. 4C, S4A). In the case of failed extrusion, the majority of cells showed defects in both basal *de novo* DJ formation and apical constriction (black in Fig. 4C). The failure of *de novo* DJ formation after desmoplakin depletion supports the molecular clamp model of desmosome function to hold cell-cell junction together in early adhesions that was described previously [29]. We defined an apical constriction defect as a failure in the closure of the apical section of the cell within two hours. Occasionally, this failure was associated with a lack of persistent constriction resulting in slight expansion of cell area after the initial constriction (Fig. S4C). Moreover, there was a lack of junctional actin accumulation at the interface between apoptotic and neighboring cells in desmoplakin-depleted monolayers compared to controls (blue arrowhead in Fig. 4A, S4B-C). A fraction of the failed extrusion cases showed only the apical constriction defect (gray in Fig. 4C, and Fig. S4D) and these defects could be explained by the effects of DJs on desmosome-junctional actin interactions [12] and nearby cytokeratin 8/18 filaments [30–32]. We further noticed similar defects related to cell extrusion and tissue integrity, in both less-dense and highly-dense tissues (Fig. S4E-E’). To complement our DP knockdown experiments, we used a monolayer overexpressing the dominant negative form of desmoplakin (DPNTP tagged to EGFP) that prevents keratin filament linkage to the desmosomal cadherins [27, 33]. Extrusion defects, including junctional tearing and delayed extrusion were observed (Fig. S4F-F”, Movie S11), which were similar to those observed in DP KD monolayers. We reasoned that lower contractility in the DP KD and DPNTP monolayers, in part, lead to such extrusion defects. To test this further, we supplemented the DP KD monolayers with the RhoA activator, CN03, before the induction of extrusion and found that the defective extrusion phenotype was rescued (Fig. 4D-D’). To further delineate the effects of desmosome at the apoptotic-non apoptotic cell interface, we followed a mosaic approach to overexpress the desmoplakin mutant, DPNTP-FLAG-EGFP either in the dying cell or in the neighboring cells (Fig. S4G-G’). When a wildtype cell was surrounded by DPNTP cells, both junctional tearing and delayed extrusion were observed, which is analogous to the phenotypes observed in DP KD and DPNTP monolayers. However, when DPNTP was overexpressed in a dying cell surrounded by wild-type cells, we observed delayed extrusion but no tissue tearing. The differences observed in these two mosaic experiments suggest that apoptosis-induced tissue tearing depends on the pre-stress of a tissue. The aberrant, desmosome-depleted junctions between dying and non-dying cell are unable to bear the tissue stress, when they are surrounded by control tissue displaying higher contractility and stress. On the contrary, the junctions are capable of bearing tissue stress when they are surrounded by DP-depleted tissue displaying lower contractility. Taken together, we conclude that desmosomes play a critical role in in the maintenance of tissue contractility to support efficient extrusion processes.

**Fig 4:**
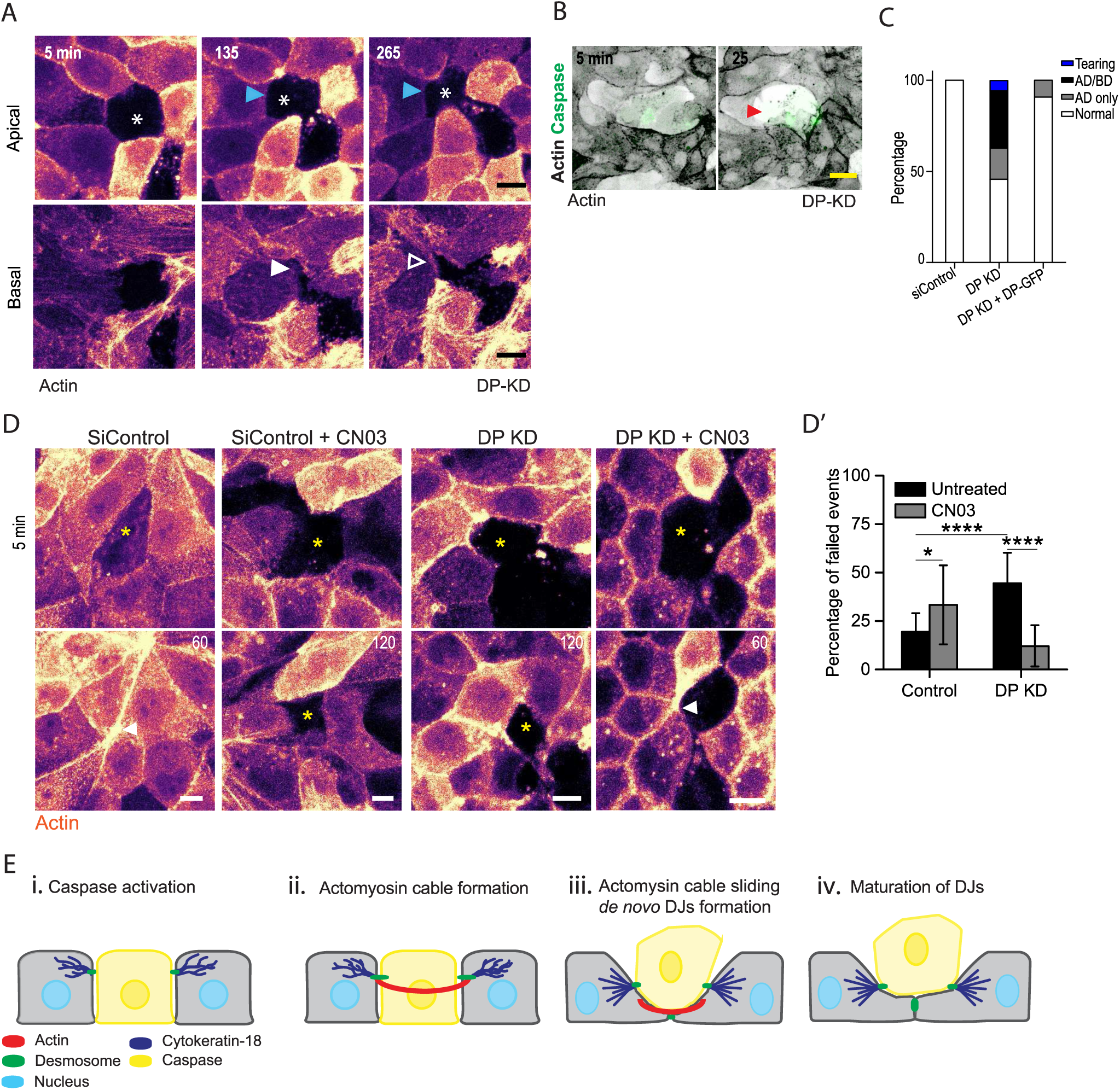
Desmosome depletion compromises apoptotic cell extrusion. A) Confocal images showing apoptotic cell extrusion under DP-depleted conditions at apical (top panel) and basal (bottom panel) sections of the cell. Asterisks and white arrowheads denote the defects in apical constriction and *de novo* cell-cell junction formation at the basal section, respectively. Blue arrowhead shows actin at the interface. B) Confocal images showing the junctional tearing phenotype at the apoptotic cell interface in DP-depleted monolayers. Red arrowhead highlights the gap in tissue. C) The percentage fraction of extrusion events in siControl control, DP-depleted and rescue conditions, including AD/BD (Apical and Basal Defects), AD only (Apical Defect only) and Normal. ‘Normal’ denotes a successful extrusion event following apical constriction, lamellipodia based movement, and *de novo* cell-cell junction formation (n=70 extrusion events from 6 independent experiments for both siControl and DP KD conditions, n=11 from 3 independent experiments for rescue condition). D) Confocal images showing representative phenotypes of siControl and DP-depleted monolayers pre-treated with CN-03. Yellow asterisks and white arrowheads denote the dying cell and completed extrusion site respectively. D’) Graph showing percentage of failed extrusion events in siControl and DP KD monolayers with/ without CN-03 pre-treatment (n=50 (siControl) and 47 (DP KD) from 4 independent experiments respectively). E) An illustration showing the interplay between desmosomes, cytokeratin-18, and actin during apoptotic cell extrusion. The data from D’ represents mean ± SEM. Two way ANOVA was used for statistical analysis of D’. *: p<0.05, ****: p<0.0001. Scale bars, 10 μm.

In this study, we reported the importance of desmosomal junctions (DJs) in mediating apoptotic cell extrusion. In contrast to E-cadherins, pre-existing DJs that originally link apoptotic cell and neighboring non-dying cells stay intact, even during the formation of *de novo* DJs between non-dying cells beneath the apoptotic cell. These observations suggest that there are two DJs at the late phase of apoptotic cell extrusion, and that mechanical coupling of the tissue is maintained throughout cell extrusion. When DJs were compromised by depletion of desmoplakin or overexpression of desmoplakin mutants, cell extrusion was compromised, likely as a consequence of a defect in apical constriction and/or defective *de novo* DJ formation (Fig. 4E). This supports the idea that DJs influence actomyosin contractility, which was subsequently confirmed by laser ablation experiments. Although the mechanism of how desmosomes influence actomyosin contractility is still not fully understood, we speculate that this could be through a mechanical and/or a biochemical link between the two cellular components.

The close proximity of DJs, associated intermediate filaments, and the actomyosin cable may mechanically link the actomyosin cable and desmosomes, either with or without cytoskeletal cross-linkers [12, 34]. A recent study reported a physical coupling between desmoglein 1 and actin via cortactin during skin differentiation, which in turn increases junctional tension in the tissue [12]. We speculate that desmoglein 2 may take part in similar mechanisms, coupling actomyosin with DJs during processes such as extrusion. This coupling is supported by our data showing that the shape of the DJs became more tortuous after the actomyosin cable departs from them during cell extrusion. Recent reports have showed how intermediate filaments that anchor at DJs regulate the contractility in proximate actin structures without a direct mechanical link but through biochemical signaling. For instance, intermediate filaments directly sequester myosin [30] and regulatory elements [31], as well as indirectly control the microtubule networks that harbor Rho GEF’s [32]. As extrusion is known to occur in multiple settings, including physiological processes, developmental events, oncogenesis, and bacterial pathogenesis [6, 35, 36], we further speculate that the defective junctions we observed in desmoplakin-depleted tissues affects actomyosin contractility and/or apoptotic cell extrusion and could lead to pathological conditions. It will be interesting to study if such extrusion defects are found in diseases and whether known mutations in both desmosomes and cytokeratin components can exacerbate diseases such as inflammatory bowel disease (IBD) and cardiomyopathies [37]. In conclusion, we showed that the coupling of the interface between dying and non-dying cells through desmosomes is required for maintaining epithelial sheet integrity during apoptotic cell extrusion.

## Author contributions

All authors designed the research. M.T. and Y.T. wrote the manuscript. M.T. performed all the experiments and analyses. Y.T. oversaw the project. All authors discussed the results and commented on the manuscript.

## Acknowledgements

This work is supported by Singapore Ministry of Education Tier 2 grant (MOE2015-T2-1-116 to Y.T.), and USPC-NUS collaborative program (to Y.T and B.L.), the Agence Nationale de la Recherche ‘‘MechanoAdipo’’ [ANR-17-CE13-0012]) and the “Labex Who Am I?” (ANR-11-LABX-0071). We are grateful to Pakorn Kanchanawong for sharing the Michael Davidson collection of plasmids, David. N. Garrod for sharing the Desmoglein 2 plasmids, James W. Nelson for sharing the MDCK Lifeact Ruby and mCherry E-Cadherin cell lines. The authors would like to thank Ong Hui Ting for discussions and help on various aspects in image processing. The authors would like to thank the MBI science communication core’s Andrew Wong and Sruthi Jagannathan for editing the manuscript and Melanie Lee for the illustrations. We also thank Virgile Viasnoff, GV Shivashankar, and members in the Toyama lab for helpful discussions.

## Supplementary figure captions

**Figure S1:**
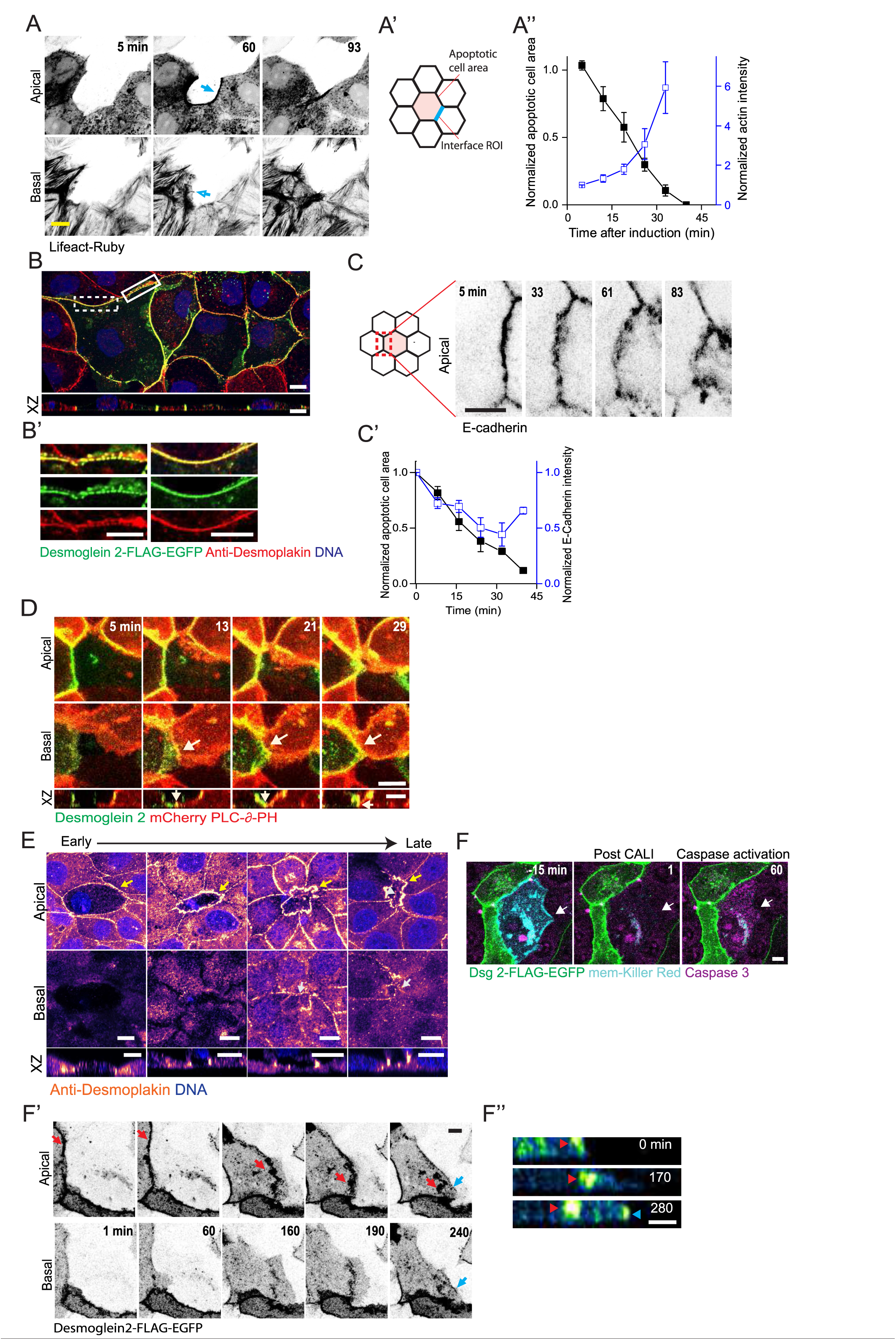
Desmosomal junctions at the interface during apoptotic extrusion. A) Confocal images showing actin (Lifeact-Ruby) dynamics during apoptotic cell extrusion in both apical (top panel) and basal (bottom panel) sections of the neighboring cell. Blue-filled and open arrows show the actomyosin cable (top panel) and the lamellipodia (bottom panel) respectively. A’) Illustration showing the definition of interface ROI during extrusion. A’’) Graphs plotting normalized apoptotic cell area (black) and actin (blue) intensity at the dying cell interface during extrusion (n=5 independent experiments). B) Confocal images showing the localization of overexpressed Dsg-FLAG-EGFP (green) and endogenous desmoplakin (red) in MDCK monolayers (n=3 independent staining experiments). B’) Magnified images of the white-filled and dotted boxes from B showing co-existence of dotted and continuous junctional clusters in the cell. C) Confocal images showing mCherry-E-cadherin dynamics at the interface during extrusion. C’) Graphs plotting normalized apoptotic cell area (black) and E-Cadherin (blue) intensity during constriction (n=5 independent experiments). D) Confocal images showing desmoglein 2 and membrane (mCherry-delta-PH) dynamics in both apical (top panel) and basal (bottom panel) sections of the cell. White arrows (bottom panel) highlights formation of *de novo* junctions (n=8 independent experiments). E) Confocal images of endogenous desmosomes in the apical (top panel) and basal (bottom panel) sections during different phases of cell extrusion. Yellow and white arrows indicate pre-existing and *de novo* junctions, respectively (n=182 extrusion events from 7 independent experiments). F) Confocal images showing desmosome (green), killer red (mem-Killer red, cyan) and caspase 3 (magenta) before and after photo-irradiation. White arrow denotes extent of the dying cell. F’) Confocal images showing desmosome dynamics (black) in the apical (top panel) and basal (bottom panel) sections of the neighboring cell. Red and blue arrows highlight pre-existing and de novo junctions respectively. F’’) Transverse view of a neighboring cell showing DJ dynamics after apoptosis induction through Killer Red. Red and blue arrows indicate pre-existing and *de novo* junctions respectively (n=4 independent experiments). The data from A’’ and C’ represent mean ±SEM. Scale bars, 10 μm except C (5 μm).

**Figure S2:**
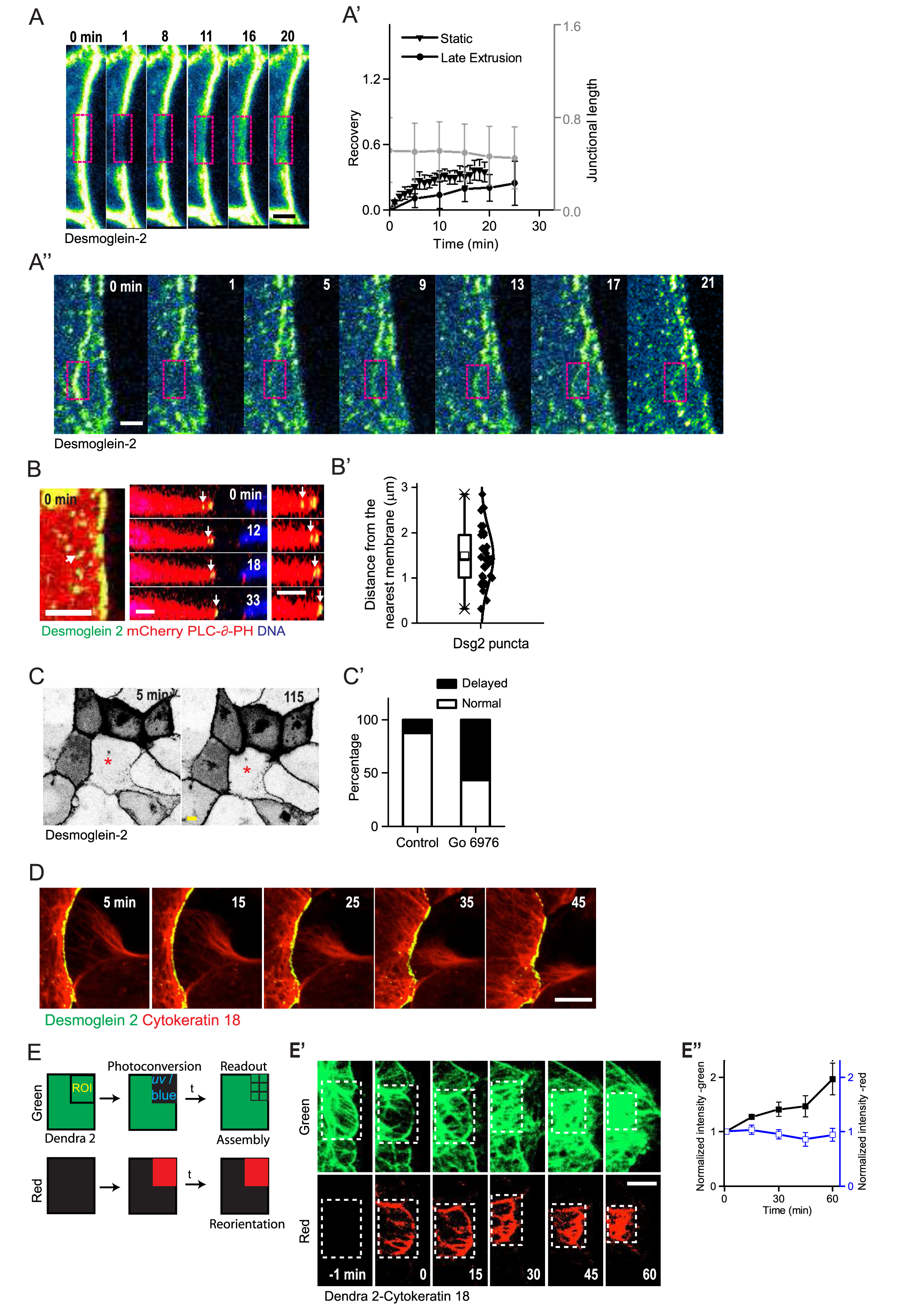
Desmosome and Cytokeratin-18 dynamics during cell extrusion. A) Confocal images showing photo-bleaching (ROI: magenta box) and recovery of static (A) and late extruding junctions (A’’) respectively. A’) Graph plotting recovery curves of late extruding junctions (black circles) and their corresponding junction lengths (gray) (n=5 and 4 independent experiments respectively). The recovery of static junctions is also plotted for comparison. B) Confocal image showing non-junctional Dsg2 puncta (green) and membrane (red) during constriction. Transverse view showing movement of puncta (white arrow) during constriction. B’) Graph plotting distance of the puncta from the nearest membrane (n=30 puncta from 5 independent experiments) C) Confocal images showing delayed extrusion following pre-treatment with Go6976. Red asterisks denote the dying cell. C’) Graph showing percentage fraction of extrusion phenotypes in control and Go6976-treated monolayers, including delayed and normal extrusion (n=8 (control) and 30 (Go6979 treated) from 3 independent experiments respectively). D) Time lapse images showing the remodeling of Cytokeratin-18 filaments through the DJ (n=6 independent experiments). E) Illustration showing the scheme of photoconversion experiment in the red (top panel) and green (bottom panel) channels respectively. E’) Time lapse images of Dendra2-Cytokeratin-18 at the apical section of the cell during extrusion in both green and red channels, representing before and after photoconversion. White dotted rectangles indicate the extent of the photoconverted region. E’’) Graph plotting normalized Cytokeratin-18 intensities of green and red channels respectively (n=4 independent experiments). The data from A’ and E’’ represent mean ±SEM and B’ represents mean ±SD. Scale bars, 5 μm except D (10 μm) and A(3 μm).

**Figure S3:**
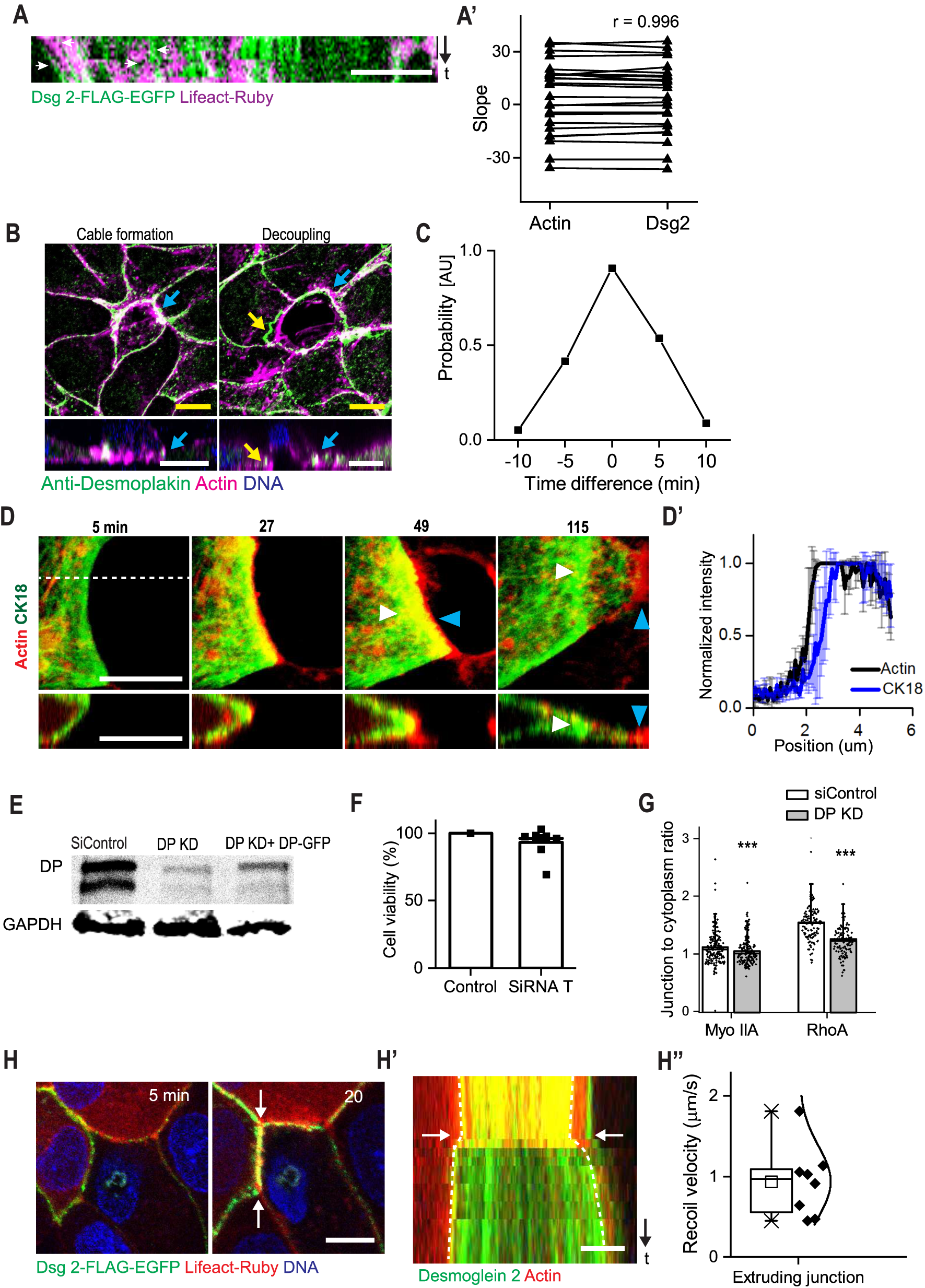
Desmosome and junctional actomyosin dynamics during extrusion. A) Kymograph showing desmosome (green) and actin (magenta) puncta movement during constriction. White arrows denote desmosome and actin puncta-associated tracks. A’) Graph plotting slope of desmosome-actin puncta pairs (n=20 puncta pairs from 5 independent experiments). B) Confocal images showing immunostained desmosome (green) and actin (magenta) during different phases of extrusion, including proximate localization of actin and desmosome (blue arrow) and detachment of actin cable from desmosome (yellow arrow) respectively (n=170 extrusion events from 4 independent experiments). C) Graph plotting the probability of time difference between actin detachment at the junctional node as compared to the middle of the junction (n=10 junctional nodes). D) Confocal images showing the temporal dynamics of mEmerald-Cytokeratin-18 (green) and Lifeact-Ruby (actin, red) during constriction. The white and blue arrowheads denote Cytokeratin 18 and actin at different phases of extrusion respectively. The white dotted line was used to generate the transverse view. D’) Line intensity profile of the dying cell interface showing proximate localization of actin (black) and keratin (blue) (n=4 independent experiments). E) Western blot images showing relative desmoplakin protein levels in siControl, DP-depleted, and rescue (DP-GFP transfected into DP depleted condition) monolayers, with GAPDH as loading control (n=3). F) Graph plotting cell viability after SiRNA transfection using Alamar blue assay (n=9 independent experiments). G) Graph showing junction-to-cytoplasm ratio of myosin IIA and RhoA in siControl and DP-depleted monolayers (n=atleast 150 junctions for each condition from 4 independent experiments). H) Confocal images showing desmosome (green) and actin (red) localization before laser ablation of the actin cable. H’) Kymograph (line along white arrows from 3H) showing recoil of desmosome (green) upon ablation of the actin cable (red). White arrows indicate time frame of ablation (n=8 independent experiments). The data from D’, F, G and H’’ represent mean ±SEM. Statistical test of G was performed using unpaired t test. *** denotes P < 0.001, respectively. Scale bars, 10 μm except H’ (5 μm).

**Figure S4:**
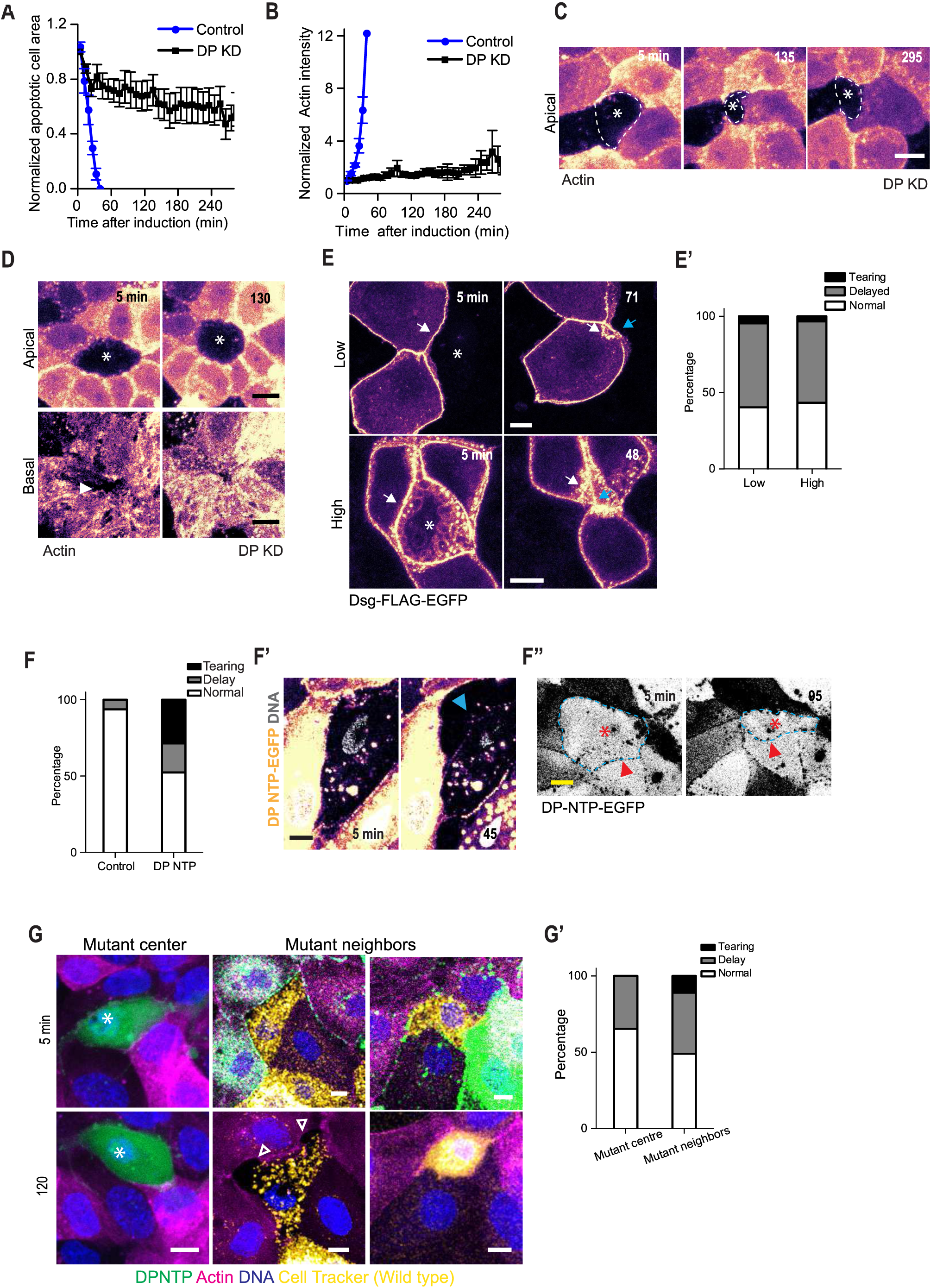
Extrusion defects on desmosome depletion. A) Graph showing the change in apoptotic cell area over time during extrusion in both siControl (blue) and DP-depleted (black) monolayers. B) Graph showing normalized actin intensity at the interface during extrusion in both siControl (blue) and DP-depleted (black) monolayers. The intensity was normalized to the corresponding value before apoptosis induction (n=6 independent experiments). C) Confocal images of actin (Lifeact-ruby) in DP-depleted monolayers showing relaxation of tissue after progressive constriction during apoptotic cell extrusion. White asterisk denotes the dying cell. D) Confocal images of actin (Lifeact-ruby) in DP-depleted monolayers showing defects in apical constriction only. The white arrowhead shows the basal area without lamellipodial crawling. Asterisk denotes defective apical constriction. E) Confocal images showing desmosome (orange) dynamics at high and low cell densities. White and blue arrows indicate pre-existing and *de novo* junctions respectively. White asterisk denotes the dying cell (n=4 independent experiments for each condition). E’) Graph showing percentage fraction of defective extrusion phenotypes under DP-depleted conditions in high and low densities, including junctional tearing, delayed constriction and normal extrusion (n=30 (high) and 42 (low) from 3 independent experiments). F) Graph showing percentage fraction of defective extrusion phenotype in wildtype and DP NTP overexpressing monolayers respectively. F’) Confocal images showing junctional tearing (blue arrowhead) at the dying cell-neighboring cell interface during extrusion. F’’) Confocal images showing delayed extrusion phenotype during the overexpression of DP-NTP (n=16 (control) and 42 (DP-NTP) from 4 independent experiments). Red asterisk denotes the dying cell. Blue dotted line and red arrowhead show extent of the cell. G) Confocal images showing extrusion defects during mosaic expression of DP-NTP mutant (green) either in dying cell (center) or neighboring cells. White arrowheads indicate junctional tearing at the dying cell-neighbor interface. White asterisks denote defective apical constriction. G’) Graph showing percentage fraction of extrusion phenotypes during mosaic expression of DP-NTP mutants, either in the dying cell (center) or the neighboring cells (n= 45 (mutant center) and 46 (mutant neighbors) from 4 independent experiments). The data from A and B represent mean ±SEM. Scale bars, 10 μm.

## Supplemental Experimental Procedures

### Cell lines, Plasmids, Si RNA and antibodies

The plasmids, mEmerald-Cytokeratin 18 (Michael Davidson collection), Desmoglein-2-Michael Sheetz), mem-killer red (Addgene,45761), DP NTP-FLAG-EGFP (subcloned from Addgene [1]), DP-GFP (Addgene,[2]) and Dendra-2-cytokeratin 18 (Michael Davidson collection) were used in this study. MDCK stable lines overexpressing Myosin Light Chain-GFP, Lifeact Ruby, mCherry-E-cadherin were kindly provided by Prof. James Nelson. The custom smartpool siRNA against Desmoplakin (XM_545329, 5’GGAAGAAGCTGGAGGACGA3’ , 5’AGGCGGAGCTGGACGGAAA3’, 5’CTACAGAACTGCTCGGATT3’, 5’GAAGATGAGAAGCGAAGAA3’) and siControl (5’UGGUUUACAUGUCGACUAA3’) were procured from Dharmacon Inc. Anti-Cytokeratin 18 (ab668, Abcam), Anti-desmoplakin (sc-33555, Santa Cruz), Anti-Myosin 2A (8064, Sigma), and Phalloidin (Invitrogen), Anti-RhoA (26C4, Santa Cruz) were used in different experiments of this study.

### Cell culture and transfection

MDCK was maintained in high glucose DMEM containing Penicillin-Streptomycin (50 U/ml), 10 % FCS and geniticin (for maintenance of stable lines). MDCK cells (10^5^) were transiently transfected with mEmerald-Cytokeratin-18 plasmid using Lipofectamine 2000 (1:2). Cells and Lip-DNA complex were mixed and seeded into a glass-bottomed dish (Iwaki, 3931-035). After 48 hours of incubation to enable sufficient incorporation of fluorescent protein into filaments, the dish was imaged using a confocal microscope (Nikon A1R MP). Dsg2-FLAG-EGFP and DP NTP-FLAG-EGFP were imaged 24 hours post transfection. The Caspase cleavable substrate (Biotium, 2 μM) and nuclear marker (Hoechst 33342, 2 μM) were added to the media prior to imaging. For visualization of membrane along with desmosome dynamics, mCherry-ò-PH was co-transfected along with Dsg2-FLAG-EGFP for 24 hours before laser-based apoptosis induction.

For mosaic experiments, wildtype cells and mutant cells were co-cultured for 24 hours before apoptosis induction. In the first case where a mutant dying cell was surrounded by normal neighbors, WT MDCK cells were transfected with DP NTP-FLAG-EGFP 24 hours prior to co-culture (mutant cells). Mutant and normal (Lifeact-ruby MDCK) cells were co-cultured in the ratio 1:10 to study extrusion dynamics under this condition. In the second case involving mutant neighbor cells and a normal center cell, the following steps were included. WT MDCK cells were labelled with Cell Tracker Deep red (Invitrogen, C34565) to enable visualization of the center cell (normal dying cell). Lifeact-Ruby MDCK cells were transfected with DPNTP-FLAG-EGFP (mutant neighbors) prior to co-culture. The normal and mutant cells were co-cultured in the ratio 1:20 followed by apoptosis induction and time lapse imaging to monitor extrusion dynamics.

### SiRNA mediated knockdown and validation

For siRNA transfection, 10^5^ cells were subjected to 50 nmoles of resuspended Desmoplakin smartpool siRNA and Dharmafect reagent. Desmoplakin knockdown was obtained 24 hours respectively post transfection and validated by western blotting. Rescue experiments were performed by transfection of DP KD cells with EGFP tagged Desmoplakin, 24 hours after the first siRNA transfection. For western blotting, 30 μg of denatured protein lysate containing RIPA buffer was loaded into a 4-15% Tris-glycine gel and subjected to SDS gel electrophoresis. The protein was transferred into a membrane, transferred by wet method and immunoblotted to visualize the protein bands by chemiluminescence. Cell viability after transfection was measured using the alamar blue assay (Invitrogen). 8000 SiRNA transfected cells were seeded in a 96 well plate and incubated with Alamar blue 24 hours post transfection. Control monolayer was not pre-treated with transfection mixture. Absorbance at 600nm and 570 nm were measured after 4 hours. Cell viability was calculated using the following formula.

% difference between control and siTransfected 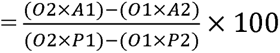

O1: Molar Extinction Coefficient of oxidized alamarBlue ^570nm^

O2: Molar Extinction Coefficient of oxidized alamarBlue^600nm^

A1: Absorbance of transfected well ^570nm^

A2: Absorbance of transfected well ^600nm^

P1: Absorbance of Control ^570nm^

P2: Absorbance of Control ^600nm^

### Laser based apoptosis induction assay

UV Laser based apoptosis induction was performed using a custom laser ablation unit interface with the NikonA1R MP microscope as described in supplemental experimental procedures [3]. The targeted nucleus was subjected to a laser power of 10-15 nW for 5 seconds to induce DNA damage, followed by the initiation of time lapse imaging.

The results from time lapse imaging during extrusion using overexpressed fluorescent protein constructs were validated by visualizing the respective endogenous components under fixed conditions. For this, WT MDCK cells were seeded into grided 35mm dishes (Ibidi, 81168). Around 10-12 cells (well-separated) of each grid square was subjected to apoptosis induction , fixed (after 60 minutes of the first apoptosis induction) and immunostained. The aforementioned grid locations were subsequently retrieved and imaged to obtain extrusion sites at different stages of extrusion.

### Killer red based apoptosis induction assay

Killer red based induction was performed by co-culture of mem-killer red expressing MDCK cells and Dsg2-FLAG-EGFP expressing MDCK cells (ratio-1:8). WT MDCK cells were transfected with mem-Killer red and Dsg2-FLAG-EGFP independently. The transfection mixture was removed after 4 hours and the monolayer was cultured in media for 24 hours. Further, they were trypsinized and co-cultured for 24 hours followed by imaging. For CALI, the green epifluorescence module of the Nikon confocal system was used for 5-7 minutes (100%). This was followed by time lapse confocal imaging to visualize caspase activation and extrusion dynamics of desmosome thereafter.

### Immunofluorescence and live imaging

Cells were fixed in either ice-cold methanol (5 minutes) or 4 % paraformaldehyde (20 minutes) in PBS. Fixed confocal imaging was performed using a spinning disk confocal microscope, 100x/1.49 objective. Validation of Dsg2-FLAG-EGFP incorporation into endogenous desmosome was performed by transfection of the plasmid into WT MDCK cells. The samples were fixed after 24 hours, immunostained (desmoplakin) and subjected to confocal imaging. Live confocal imaging was performed using Nikon A1R MP with a 60X Nikon Apo 60x/1.40 oil immersion objective lens at 37**°**C, 5% CO_2_ in a humidified incubation chamber. For high time resolution (Fig. S3A) and observation of sub-junctional puncta, time lapse imaging was performed once every 15 seconds on a single stack.

### FRAP and Photoconversion

Dsg 2-FLAG-EGFP was overexpressed in wild type MDCK cells. A rectangular ROI, 2*1 μm was defined at the interface. Bleaching was performed using the 488 laser at 20% laser power, scanning speed-8 in the NikonA1R MP microscope. FRAP was performed on junctions under three conditions-static junction (non-apoptotic), early extrusion (Bleaching 5 minutes after induction) and late extrusion (bleaching after the de novo junction formation). The Z stack was imaged for 30 minutes post bleaching. For photoconversion experiments, Dendra2-Cytokeratin18 was overexpressed in WT MDCK cells. Photoconversion of green filaments into red, was performed at an interface ROI using a 405nm laser. A rectangular ROI 2*1 μm^2^ was drawn on the interface of the neighboring cell. Photoconversion was performed using the 405 nm laser line of the Nikon A1R MP, using the following conditions: 5% laser power, scanning speed-8 and two iterations. Laser based apoptosis induction was performed following this and both red and green channels were imaged for 1 hour.

### Laser ablation

The apical sections of the cell, with highest Lifeact Ruby intensity, were used for junctional ablation. This experiment was performed using the ultra violet laser ablation system (355 nm, 300 ps pulse duration, 1 kHz repetition rate, PowerChip PNV-0150-100, team photonics) as described in [4]. The ablation parameters were optimized as 120 nW, with an exposure time of 0.3 seconds. Imaging was performed thereafter, at an interval of 2.2 seconds for a time frame of 90 seconds. Keratin filament network ablation was performed, by introducing a line cut (6-8 μm) using the same laser at 75 nW, either 5 minutes (early) or 45-60 minutes (late, after cytokeratin reorientation) after induction of apoptosis (laser-based apoptosis induction assay). For ablation of actin cable during extrusion, apoptosis was induced and time lapse imaging was set up to monitor extrusion dynamics until actin cable formation. At this stage, a line cut (1-2 μm) across the junction was performed at 100 nW and time lapse images were acquired at an interval of 2 seconds to capture recoil of desmosome junctional nodes.

### Drug treatment

Cells were pretreated with DMSO control or 1 µM Go6976 for 30 minutes followed by laser-based apoptosis induction. For RhoA hyperactivation, siControl and DP-KD monolayers were first subjected to apoptosis (laser-based apoptosis induction assay) and time-lapse imaging to evaluate extrusion dynamics. The same samples were subsequently incubated with 0.25 mM CN-03 for 2 hours at 37°C. The samples were further subjected to apoptosis induction and time lapse imaging to evaluate extrusion dynamics under hypercontractile conditions.

### Image processing and quantification

Image processing and quantification was performed using different plugins in Fiji[5].

### Junctional interface intensity, cell spread area and Z position

Quantification of cell spread area and interfacial Lifeact (/desmoglein / cytokeratin 18) intensity was computed by drawing a polgon and line ROI (5 pixel wide) respectively at the dying cell interface and stored using the ROI manager. The basal slices with actin stress fibers were excluded from all data sets to avoid interference from the basal cytoskeleton. The sum of slices projection was used for intensity measurement. Normalized intensity (intensity per unit length at t_n_/ intensity per unit length at t_1_) was calculated and plotted as a function of time. The calculation of time taken for extrusion of control and knockdown/mutants were performed similarly.

The Z position of DJ and actin during extrusion was computed by generating the transverse view using the “Reslice” function of Fiji. The XY coordinates of the centroid of the junction and actin were obtained and quantified. Position of Dsg2 puncta in relation to the nearest membrane was computed by generating a transverse section using the “Reslice” function and the distance of the puncta (green) to the nearest membrane (apical, basal and junctional edges) (red) was measured.

### FRAP and photoconversion

For quantification of FRAP, the ROI was changed manually across the time series, after taking into consideration the junctional shortening and the change in bleaching ROI thereof during extrusion. A line (width: 5 pixel) was drawn across the newly defined ROI between the junctions to measure the recovery during different stages of extrusion. Normalized intensity of the photobleached ROI (Recovery= intensity at t_n_/ pre-bleaching intensity(t_0_)) was calculated and plotted as a function of time. The photoconverted ROI was changed in the time series, w.r.t the extent of the intensity in the red channel. The intensity was measured using a rectangular ROI. The normalized intensity (normalized intensity _=_ intensity at t_n_/ intensity at t_1_) of both the red and green channels was measured and plotted. t_1_ indicates the intensity at photoconversion.

### Straightness index

Straightness index, an chord-arc ratio (SI= Junctional node displacement L_0_ / path length L) was obtained by manual segmentation of the junction at different time points using Fiji, to obtain both the junctional length as well as displacement between the junctional nodes.

### Orientation analysis

Orientation of the global Cytokeratin18 network during the progression of extrusion was computed by measuring the structure tensor of the timelapse images. This was performed using the OrientationJ plugin of Fiji [6, 7]. The images in the time series were background subtracted, and rotated parallel to the line connecting the centroids of the dying cell and the neighbor at t_1_. The orientation plots and the corresponding angle distribution were obtained. The color survey was generated by setting the three parameters; hue, saturation and brightness to orientation, original image and constant respectively. The orientation angles were obtained and plotted as a distribution.

### Quantification of junctional Actin, Myosin IIA and RhoA intensity

Junctional actin was computed using a custom Matlab algorithm. The algorithm performed local thresholding and segmented the junctions, followed by separation of the branched nodes, to enable intensity quantification of individual junctions. The junctional actin was measured by drawing a junctional line ROI (width: 5 pixel).

Junctional myosin IIA and Rho A were quantified manually using a Fiji macro, owing to the poor cytoplasm-junction contrast. The segmentation of the junctional pools of myosin IIA and Rho A were performed manually. The basal sections of the cell were removed, and the apical slices were used generate a projection (sum of slices), followed by drawing a segmented line ROI (width: 5 pixel) to measure the corresponding intensity. Normalized intensity of DP KD junction (Normalized intensity at J_KD1_= Intensity at J_KD1_/ Average intensity of control junctions) was calculated and plotted. To further compute junction to cytoplasm ratio (ratio = Average intensity of junction I_junc_/ Average intensity of cytoplasm I_cyto_), a circular cytoplasmic ROI (5 µm diameter, nucleus excluded and adjacent to the corresponding junction) was drawn on the projected image to obtain the cytoplasmic intensity inaddition to the junctional intensity obtained as described above. The intensity of DP KD associated junctions was normalized with respect to the normal junctions and plotted.

### Movement of actin and Dsg2 puncta

A kymograph was drawn manually along the junction during constriction. Tracks of Dsg2 and Lifeact-Ruby pairs were drawn manually. The slope was computed by obtaining the line coordinates and plotted.

### Actin decoupling

Time difference (ΔT) is defined as the time lag between detachment of desmosome at the middle of the junction as compared to the junctional nodes. ‘0’ indicates simultaneous decoupling. The probability density function of ΔT was computed and plotted.

### Junctional perturbation studies

An extrusion site where the dying cell is removed from the monolayer within 120 minutes is defined as ‘normal’. An extrusion site where the extruding cell remains in the epithelium after 120 minutes of apoptosis induction was defined as ‘delayed’. An extrusion site where the extruding cell loses contact with the neighbor cells was defined as ‘tearing’.

### Recoil velocity

Recoil velocity was computed by tracking the Cartesian coordinates of the junction nodes using MTrackJ plugin in Fiji [8], followed by fitting the calculated distance between nodes into a single/double exponential function. Recoil velocity was calculated as the derivative of the function using a custom made MATLAB algorithm as described in [4].

### Statistical analysis

The data indicated is mean ± SEM. The number of independent experiments performed is indicated in the figure captions. Atleast 4 independent biological replicates were performed for each experiment. Statistical tests were performed using Graphpad Prism and Origin Pro 6.0. Unpaired t test (with Welch correction) (Fig. 2D’, 3C, 3D’ S2C’, S3G) and Paired t test (Fig.3A’’) were used to test the statistical significance.

